# *Salmonella* exploits OmpR to switch essential morphogenetic peptidoglycan synthases in response to intracellular cues

**DOI:** 10.1101/2021.10.28.466273

**Authors:** David López-Escarpa, Sónia Castanheira, Francisco García-del Portillo

## Abstract

Essential peptidoglycan synthases, like penicillin binding proteins 2 and 3 (PBP2/PBP3) of *Escherichia coli*, define shape by orchestrating cell elongation and division, respectively. Despite being intensively studied as drug targets, the regulatory rules governing their production remain poorly understood. During infection, the closely related pathogen *Salmonella enterica* serovar Typhimurium downregulates PBP2/PBP3 production and replace them with alternative peptidoglycan synthases, PBP2_SAL_/PBP3_SAL_, absent in *E. coli*. The bases for such switch in morphogenetic proteins are unknown. Here, we show that the *S.* Typhimurium regulator OmpR triggers PBP2_SAL_ and PBP3_SAL_ expression responding solely to acid pH and define a shared motif present in upstream regions of the PBP2_SAL-_ and PBP3_SAL_-coding genes sufficient for such control. The elimination of PBP2/PBP3 in infection conditions is however multifactorial, requiring acidity, high osmolarity and being favoured by OmpR and the Prc protease. Remarkably, we found that *E. coli* loses the essential PBP3 required for cell division when exposed to both acidity and high osmolarity, the environmental cues encountered by intracellular *S.* Typhimurium. Therefore, OmpR played a central role in the evolution of this pathogen when co-opting the regulation of PBP2_SAL_/PBP3_SAL_ and, consequently, promoting a new morphogenetic cycle that made possible increasing progeny inside acidic eukaryotic phagosomes.

**Significance:** Some enzymes that participate in peptidoglycan metabolism are present exclusively in bacterial pathogens and modify its structure to limit immune recognition. The intracellular pathogen *Salmonella enterica* serovar Typhimurium is the only example known to date in which a “substitution” of essential peptidoglycan enzymes involved in cell division and elongation takes place during infection. The data presented here support instability of PBP3 in environments with acidity and high osmolarity as a probable selective pressure that promoted the fixation of alternative morphogenetic enzymes. This was possible due to the control that OmpR exerted over these new foreign functions. The acquisition of enzymes like PBP2_SAL_ and PBP3_SAL_ therefore represent a “quantum leap” evolutionary event in *S.* Typhimurium that made possible the colonization of acidic intracellular niches.

## Introduction

The peptidoglycan (PG), also known as murein sacculus, marked the evolution of the domain Bacteria as main component of the bacterial cell wall (Egan et al., 2020). It is remarkable among its physicochemical properties the presence of D-amino acids and its assembly as a single giant covalently-bound macromolecule covering the entire cell surface. Based on this, the synthesis and remodelling of its structure are pivotal for determining cell shape and constructing the cell division septum that separates daughter cells (Egan et al., 2017; Egan and Vollmer, 2013). The expansion of the PG along the cell surface has been extensively studied in rod-shaped bacteria like *Escherichia coli* and *Bacillus subtilis*, in which the cell elongation and division phases are clearly defined temporally and spatially (den Blaauwen et al., 2008; Errington, 2015; Rohs and Bernhardt, 2021). Both phases are commanded by monofunctional PG synthases with transpeptidase (TP) activity that crosslink stem peptides in parallel glycan chains of the sacculus (den Blaauwen et al., 2017; Szwedziak and Lowe, 2013). This reaction proceeds via recognition of the terminal D-alanine-D-alanine present in the stem peptide of the lipid II precursor molecule, which in *E. coli* is *N*-acetyl-glucosamine-*N*-acetyl-muramyl-L-alanine-D-glutamic acid-*meso*-diaminopimelic acid*-*D-alanine-D-alanine (NAG-NAM-L-Ala-D-Glu-*m*-DAP-D-Ala-D-Ala) (Typas et al., 2011).

In *E. coli,* the monofunctional PG synthases with TP activity that control cell elongation and cell division are the penicillin-binding proteins 2 and 3 (PBP2 and PBP3), respectively (Egan et al., 2017; Szwedziak and Lowe, 2013). PBP2 and PBP3 form part of multiprotein complexes named elongasome and divisome that have RodA and FtsW as cognate glycosyltransferases (GT). These PBP2-RodA and PBP3-FtsW pairs facilitate the incorporation of the lipid II precursor to the growing PG sacculus during the phases of cell elongation and division (Daitch and Goley, 2020; den Blaauwen and Luirink, 2019; Rohs and Bernhardt, 2021). Importantly, the elongasome and divisome are connected to cytoskeletal scaffolds that guide the synthetic machinery to either defined locations along the cylindrical region of the cell or the transversal septum required for division (Cho, 2015; Errington, 2015; McQuillen and Xiao, 2020). Due the important role of these two multiprotein complexes for the morphogenesis and proliferation of bacteria, PBP2 and PBP3 are essential enzymes and have been historically the target of choice for the development of effective beta-lactam antibiotics.

The genome of *Salmonella enterica* serovar Typhimurium (*S.* Typhimurium) encodes a pair of PBPs, named PBP2_SAL_ and PBP3_SAL_, which replace PBP2 and PBP3 during the infection of susceptible mice by this intracellular pathogen (Castanheira et al., 2020). PBP2_SAL_ and PBP3_SAL_ are paralogue enzymes (~67% of identity at the protein level) active in acid pH and up-regulated by *S.* Typhimurium after the invasion of eukaryotic cells (Castanheira et al., 2020, 2017). In laboratory media, PBP2_SAL_ and PBP3_SAL_ production is exacerbated in acidified nutrient media. Thus, the substitution of PBP2/PBP3 by PBP2_SAL_/PBP3_SAL_ detected *in vivo* is partially reproduced when bacteria grow in acidified media with limited amount of nutrients (Castanheira et al., 2020).

Considering this unprecedented replacement of essential morphogenetic PG enzymes when *S.* Typhimurium adapts to distinct lifestyles and the stimulation of such phenomenon by intracellular cues like acid pH and high osmolarity, we sought to dissect the underlying regulatory network responsible for this phenomenon. Here, we show that OmpR is the master regulator that triggers the production of PBP2_SAL_ and PBP3_SAL_. OmpR was first identified in *E. coli* as modulator of relative levels of major outer membrane proteins like OmpC and OmpF (Taylor et al., 1981) and, subsequently reported to be a positive regulator of the *Salmonella*-pathogenicity island-2 (SPI-2) that modulates the *S.* Typhimurium intracellular lifestyle (Feng et al., 2003; Garmendia et al., 2003; Lee et al., 2000). OmpR has also been reported to control in a positive or negative manner the expression of virulence factors in other pathogens like enterohemorrhagic *E. coli* (Wang et al., 2021), *S.* Typhi (Santander et al., 2008), or *Vibrio cholerae* (Kunkle et al., 2020). Our findings extent this view and catalogue OmpR as a key regulator of PG metabolism and morphogenesis when *S.* Typhimurium colonizes the intracellular niche of eukaryotic cells.

## Results

OmpR triggers the expression of PBP2_SAL_ and PBP3_SAL_ in response to acid pH. To adapt to the intracellular lifestyle inside acidic vacuoles, *S.* Typhimurium uses regulators like the two component regulatory systems PhoP-PhoQ, EnvZ-OmpR, PmrA-PmrB and SsrA-SsrB (Chen and Groisman, 2013; Dalebroux and Miller, 2014; Feng et al., 2003; Kenney, 2019; Pérez-Morales et al., 2017) and the transcriptional regulator SlyA (Buchmeier et al., 1997), among others. PBP2_SAL_ and PBP3_SAL_ production is up-regulated inside host cells (Castanheira et al., 2017), so we sought to determine whether any of these regulators was involved. Null alleles of the genes encoding these regulators were introduced in a *S.* Typhimurium strain that was 3xFLAG epitope-tagged in the chromosome at the *SL1344_1845* and *SL1344_1765* genes, encoding PBP2_SAL_ and PBP3_SAL_ respectively. The 3xFLAG tag was placed at the 3’ end of the respective coding sequences. Our previous studies show that PBP2_SAL_-3xFLAG and PBP3_SAL_-3xFLAG are functional since they bind beta-lactams and contribute to morphogenesis in mutants lacking PBP2 or PBP3 [(Castanheira et al., 2017) and data not shown].

Relative levels of PBP2_SAL_-3xFLAG and PBP3_SAL_-3xFLAG (hereinafter referred as PBP2_SAL_ and PBP3_SAL_) did not change drastically in the isogenic regulatory mutants except for the *ompR* mutant, which produced almost undetectable levels of both enzymes (Fig. 1A-B), and the *phoP* and *slyA* mutants with slightly lesser amounts of PBP2_SAL_ (Fig. 1A). Based on this prominent role of OmpR in controlling production of these PG synthases, we next considered a recent transcriptomic study reporting *dacC*, encoding the carboxypeptidase PBP6 that trims stem peptides of the PG, as an OmpR-regulated gene in *E. coli* and *S.* Typhimurium (Chakraborty and Kenney, 2018). Using chromosomally *dacC*::3xFLAG-tagged strains, we did not detect such regulation at the protein level with wild type and *ompR* strains having similar amounts of PBP6 (DacC) (Fig. 1C). Similarly, no changes in the amounts of the bifunctional PG synthases PBP1A and PBP1B were detected in response to acid pH (Fig. S1).

**Figure 1.**
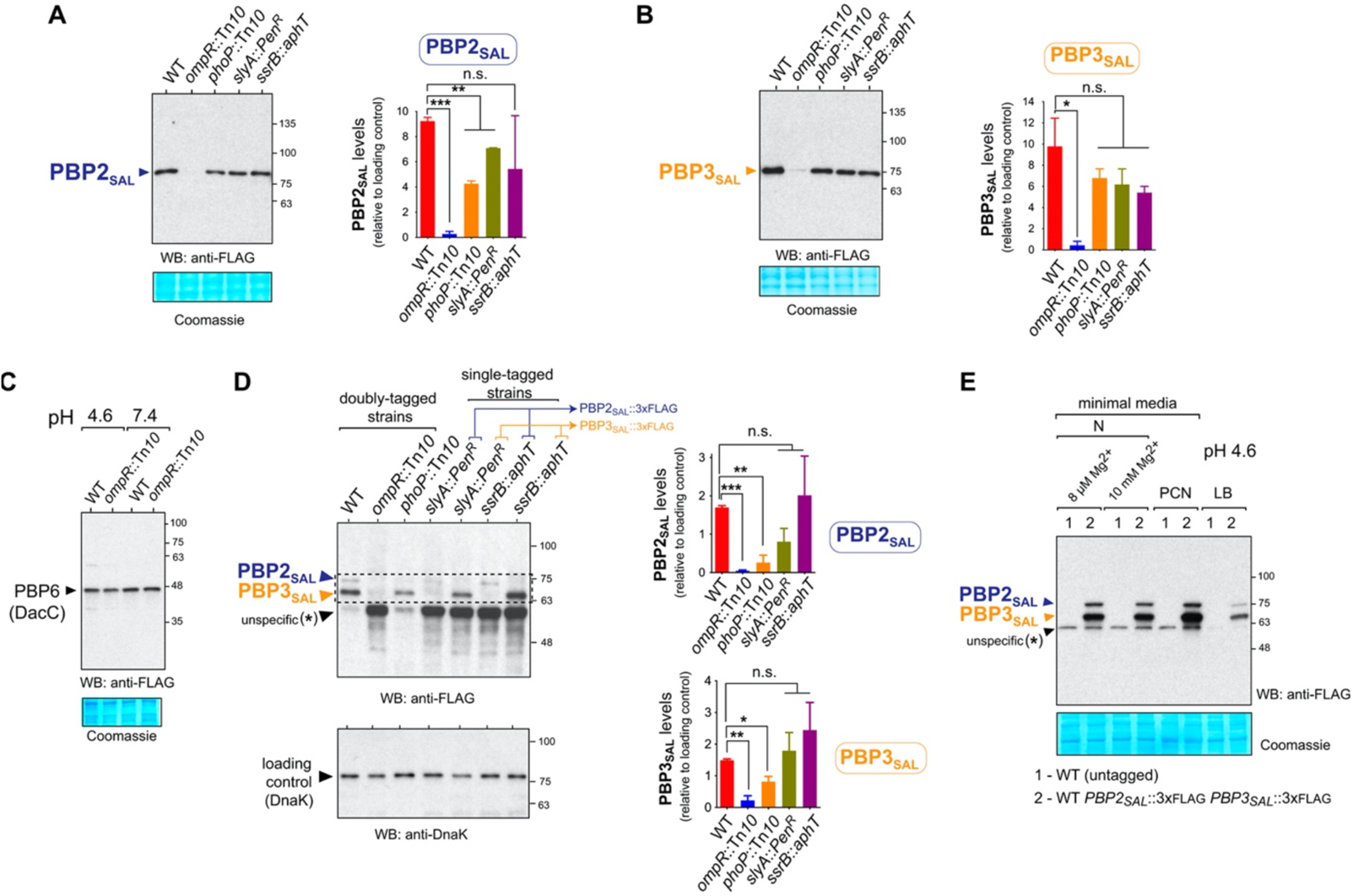
OmpR is the main regulator controlling the production of PBP2SAL and PBP3SAL in *S.* Typhimurium. (A) Immunodetection of PBP2SAL in total protein extracts of isogenic tagged *S.* Typhimurium strains bearing a PBP2SAL-3xFLAG allele in its native chromosomal location and mutations in the indicated regulators (OmpR, PhoP, SlyA and SsrB) when growing in LB medium pH 4.6. Shown are also the loading control (Coomassie staining) and the quantification data of the PBP2SAL protein levels; (B) Same as for (A) but for isogenic strains bearing a PBP3SAL-3xFLAG allele in its native chromosomal location; (C) Unlike PBP2SAL and PBP3SAL, the levels of the carboxypeptidase PBP6 (DacC) involved in PG metabolism, remain unaltered in the absence of OmpR. For this assay, isogenic strains bearing a *dacC*::3xFLAG allele in its native chromosomal location were grown in minimal N medium at the indicated pH (4.6 or 7.4); (D) Levels of PBP2SAL and PBP3SAL produced by intracellular *S.* Typhimurium at 8 h post-infection of NRK-49F rat fibroblasts. Isogenic wild-type and mutant strains lacking the indicated regulators and either doubly or singly-tagged with the PBP2SAL-3xFLAG and PBP3SAL-3xFLAG alleles in their respective chromosomal locations, were used. DnaK was used as loading control. Data shown in panels A-D are means and standard deviations from a minimum of two independent experiments and statistically analyzed by unpaired parametric t-test. *, *P* = 0.01 to 0.05; **, *P* = 0.001 to 0.01; ***, *P* = 0.0001 to 0.001; n.s., not significant; (E) Detection of an unspecific band in both untagged and tagged *S.* Typhimurium strains in immunoassays with the anti-FLAG antibody when using electrophoresis system based on precast 4-20% gradient gels required to separate and resolve the positions of PBP2SAL and PBP3SAL (see Materials and Methods). Total protein extracts were prepared from bacteria grown in the indicated minimal (N, PCN), and nutrient rich (LB) media at pH 4.6.

**Figure S1.**
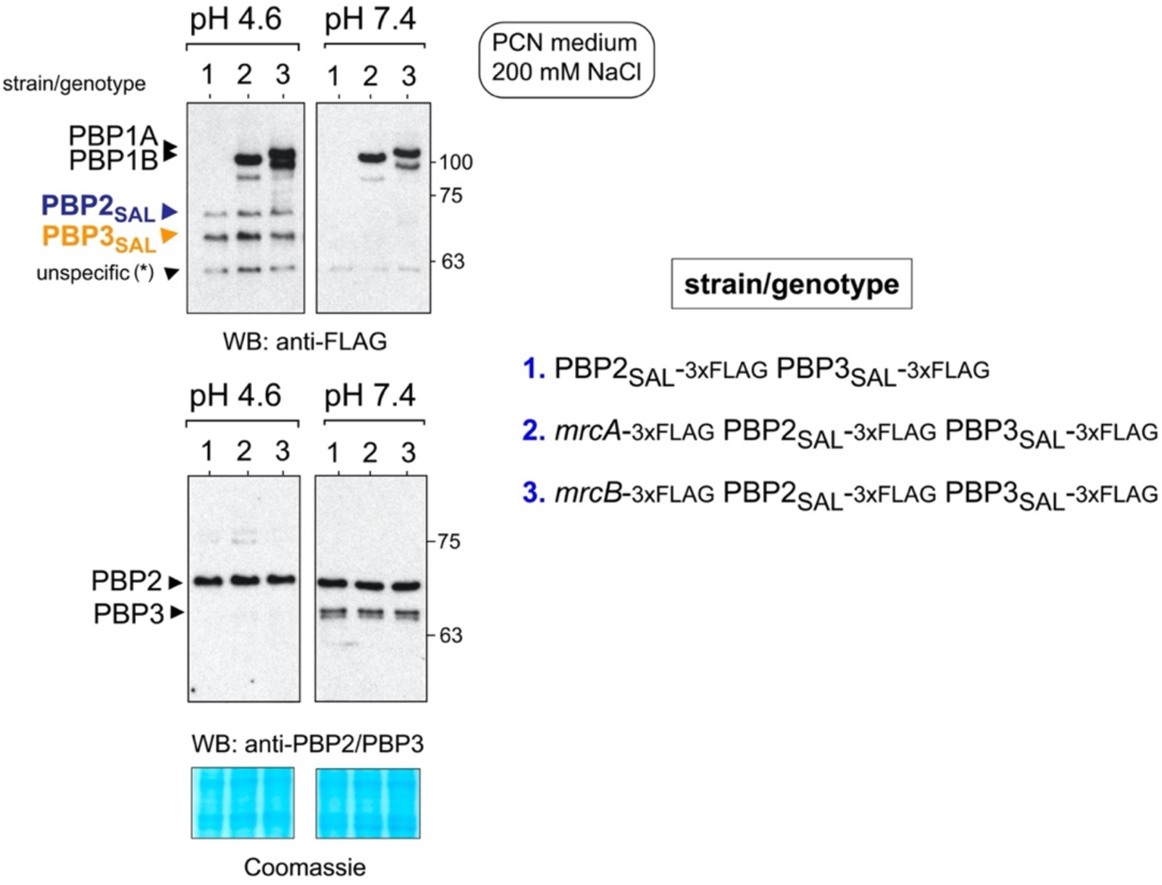
The production of the bifunctional PG synthases PBP1A and PBP1B is not altered by pH in *S.* Typhimurium. Shown are western blots of total protein extracts of the indicated chromosomally tagged *S.* Typhimurium strains grown in PCN medium 200 mM NaCl at pH values of 4.6 and 7.4. The Coomassie stained membrane is shown as loading control. Note that, unlike PBP1A and PBP1B, the PBP2SAL and PBP3SAL enzymes are produced exclusively in acid pH, a condition in which PBP3 is not detected. Data representative of a total of three independent repetitions.

We next determined whether OmpR regulates PBP2_SAL_ and PBP3_SAL_ in intracellular *S.* Typhimurium after invasion of eukaryotic cells (Fig. 1A-B). This was confirmed in cultured fibroblasts since the *ompR* mutant did not produce PBP2_SAL_ or PBP3_SAL_ (Fig. 1C). A substantial decrease of PBP2_SAL_ and PBP3_SAL_ (20%, 50% of wild type values, respectively) was observed for the *phoP* mutant (Fig. 1D). Not statistically significant differences in PBP2_SAL_ and PBP3_SAL_ levels were however found for intracellular bacteria lacking either SlyA or SsrB (Fig. 1D). In some instances, like the analysis performed with extracts prepared from intracellular bacteria, the immunodetection of PBP2_SAL_-3xFLAG and PBP3_SAL_-3xFLAG was accompanied by the appearance of a prominent unspecific band (see Fig. 1D). This band was unrelated to PBP2_SAL_-3xFLAG and PBP3_SAL_-3xFLAG as was also detected in untagged wild type bacteria (Fig. 1E). Curiously, this unspecific band was noticeable in protein extracts resolved in precast 4-20% gradient gels. This type of gel was however the most useful for increasing the difference in the electrophoretic mobilities of PBP2_SAL_-3xFLAG and PBP3_SAL_-3xFLAG.

We also examined whether the alternative sigma factors RpoS and RpoE, used by S. Typhimurium to grow and survive inside host cells (Cano et al., 2001; Chen et al., 1996; Osborne and Coombes, 2009), could regulate PBP2_SAL_ and PBP3_SAL_ production. Mutants lacking these sigma factors produced however wild type levels of both enzymes (Fig. S2). Taken together, these data indicated that OmpR is the master regulator that controls the production of PBP2_SAL_ and PBP3_SAL_ in both acidified laboratory media and inside eukaryotic cells.

**Figure 2.**
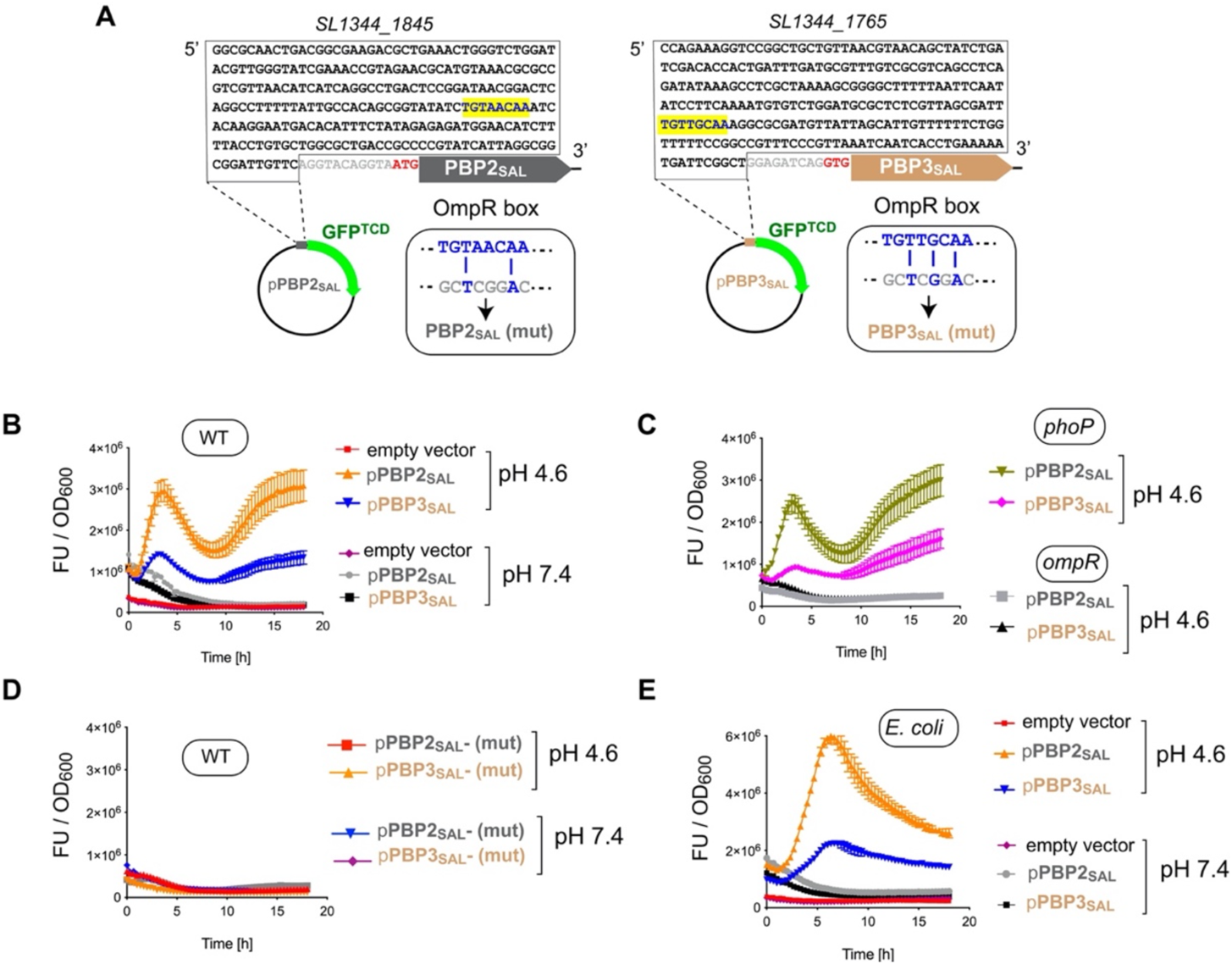
The promoter regions of the *S.* Typhimurium genes encoding PBP2SAL and PBP3SAL harbor sequences recognized by OmpR. (A) Upstream regions of the genes encoding PBP2SAL and PBP3SAL that were cloned in the promoter-less reporter plasmid expressing the GFP^TCD^ variant. In yellow are indicated the OmpR boxes analysed and the insets depict the mutated versions (mut) generated in each of the two regions; (B-E) Activity of the PBP2SAL and PBP3SAL promoters in the following strains and growth conditions: (B) wild type *S.* Typhimurium, PCN medium 200 mM NaCl at neutral and acid pH (7.4 and 4.6, respectively); (C) isogenic *S.* Typhimurium mutants lacking either PhoP or OmpR grown in PCN medium 200 mM NaCl pH 4.6; (D) wild type *S.* Typhimurium bearing reporter constructed based on the mutated (mut) versions of each promoter; and, (E) wild type *E. coli* expressing the reporter constructs based on the *S.* Typhimurium PBP2SAL and PBP3SAL promoters. Data were collected automatically during 18 h in a TECAN plate reader and are represented as the mean and standard deviation from three technical replicates from a representative experiment of a total of three biological replicates. FU, fluorescent units.

**Figure S2.**
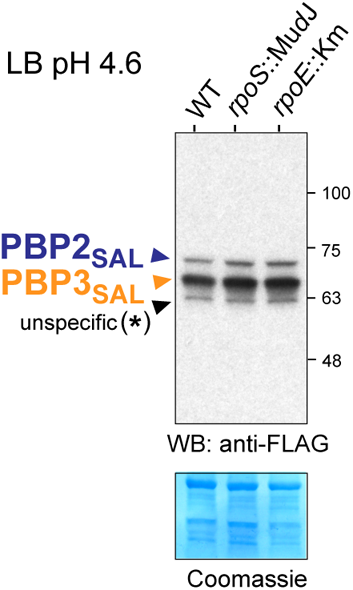
The alternative sigma factors RpoS and RpoE are not required for the production of PBP2SAL or PBP3SAL in response to acid pH. Shown are total protein extract of isogenic *S.* Typhimurium wild type and mutant strains defective in RpoS or RpoE grown in LB medium at acid pH of 4.6. The Coomassie stained membrane is shown as loading control. Data representative of a total of three independent repetitions.

OmpR recognizes sequence motifs located in the upstream region of the genes encoding PBP2_SAL_ and PBP3_SAL_. OmpR binds to target genes by recognizing sequence motifs in the upstream regulatory regions by mechanisms involving relaxation of supercoiled DNA and release of repression mediated by nucleoid binding proteins like H-NS and Fis (Banda et al., 2019; Cameron and Dorman, 2012). ChiP-Seq studies performed in *S.* Typhimurium and *S.* Typhi reported TGT(A/T)ACA(A/T) as consensus binding site for OmpR (Perkins et al., 2013). We searched *in silico* for putative OmpR binding sites in 250 nt upstream regions of *SL1344_1845* and *SL1344_1765*, the genes of *S.* Typhimurium strain SL1344 that encode PBP2_SAL_ and PBP3_SAL_ respectively. Each of these regions has a putative OmpR-binding site matching the consensus: TGTAACAA in *SL1344_1845* (PBP2_SAL_) and TGTTGCAA in *SL1344_1765* (PBP3_SAL_) (Fig. 2A). The distance of these boxes relative to the start codon is 104 nt for *SL1344_1845* and 91 nt for *SL1344_1765* (Fig. 2A).

We cloned these 250 nt upstream regions harbouring the putative OmpR binding sites in a promoter-less vector bearing the *gfp*^TCD^ gene redesigned to reduce H-NS transcriptional silencing and having improved translation rate (Corcoran et al., 2010). The upstream regions of PBP2_SAL_ and PBP3_SAL_ triggered GFP^TCD^ expression responding to acid pH and in an OmpR dependent manner albeit independently of PhoP (Fig. 2B-C). We next replaced the putative OmpR boxes identified in the PBP2_SAL_ and PBP3_SAL_ upstream regions by a non-functional GCTCGGAC mutated sequence (Fig. 2A), previously reported to abolish OmpR-mediated expression in the *S.* Typhi *tviA* gene (Perkins et al., 2013). This mutated sequence resulted in lack of GFP^TCD^ expression (Fig. 2D), therefore indicating that the two sequence motifs identified *in silico* in the upstream regions of *SL1344_1845* (PBP2_SAL_) and *SL1344_1765* (PBP3_SAL_) are *bona fide* OmpR binding sites. The transformation of *E. coli* with the reporter plasmids bearing the wild type OmpR boxes of the PBP2_SAL_ and PBP3_SAL_ upstream regions led to a similar expression pattern as observed in *S.* Typhimurium, strictly dependent on acid pH (Fig. 2E). Altogether, these data showed that *S.* Typhimurium OmpR triggers expression of PBP2_SAL_- and PBP3_SAL_-coding genes recognizing specific sequences in their upstream binding sites and, that this regulation can be recapitulated in *E.coli*, therefore ruling out the need of *Salmonella*-specific co-regulators.

OmpR controls the production of PBP2_SAL_ and PBP3_SAL_ responding exclusively to acid pH. The EnvZ-OmpR two component system senses acid and osmotic stress to regulate gene expression (Chakraborty and Kenney, 2018). We used minimal defined media to determine whether, in addition to acid pH, other signals such as high osmolarity are required for PBP2_SAL_ and PBP3_SAL_ expression. PBP2_SAL_ and PBP3_SAL_ levels were monitored in PCN minimal medium, in the presence/absence of OmpR, pH values 4.6 and 7.4 and salt concentrations of 0 and 200 mM NaCl. This high amount of salt triggers in *E. coli* K-12 maximal changes in expression of outer membrane proteins further shown to be regulated by OmpR (Alphen and Lugtenberg, 1977; Verhoef et al., 1979). Our control tests showed that 200 mM NaCl does not affect viability in *S.* Typhimurium (see below).

The assays in PCN minimal medium showed that acid pH is the only signal required for triggering PBP2_SAL_ and PBP3_SAL_ production in an OmpR-dependent manner (Fig. 3A). Thus, high salt concentration (200 mM NaCl) did not increase notoriously the levels of any of the two enzymes (Fig. 3A). Remarkably, we noted that PBP2 and PBP3 levels, which were monitored in parallel in the same samples, decreased drastically when bacteria were exposed to both signals, acid pH of 4.6 and 200 mM NaCl (Fig. 3A). *S.* Typhimurium OmpR contributes minimally to the substantial decrease of PBP2 and PBP3 in pH 4.6 200 mM NaCl, as only PBP3 levels increased slightly in the absence of this regulator (Fig. 3A).

**Figure 3.**
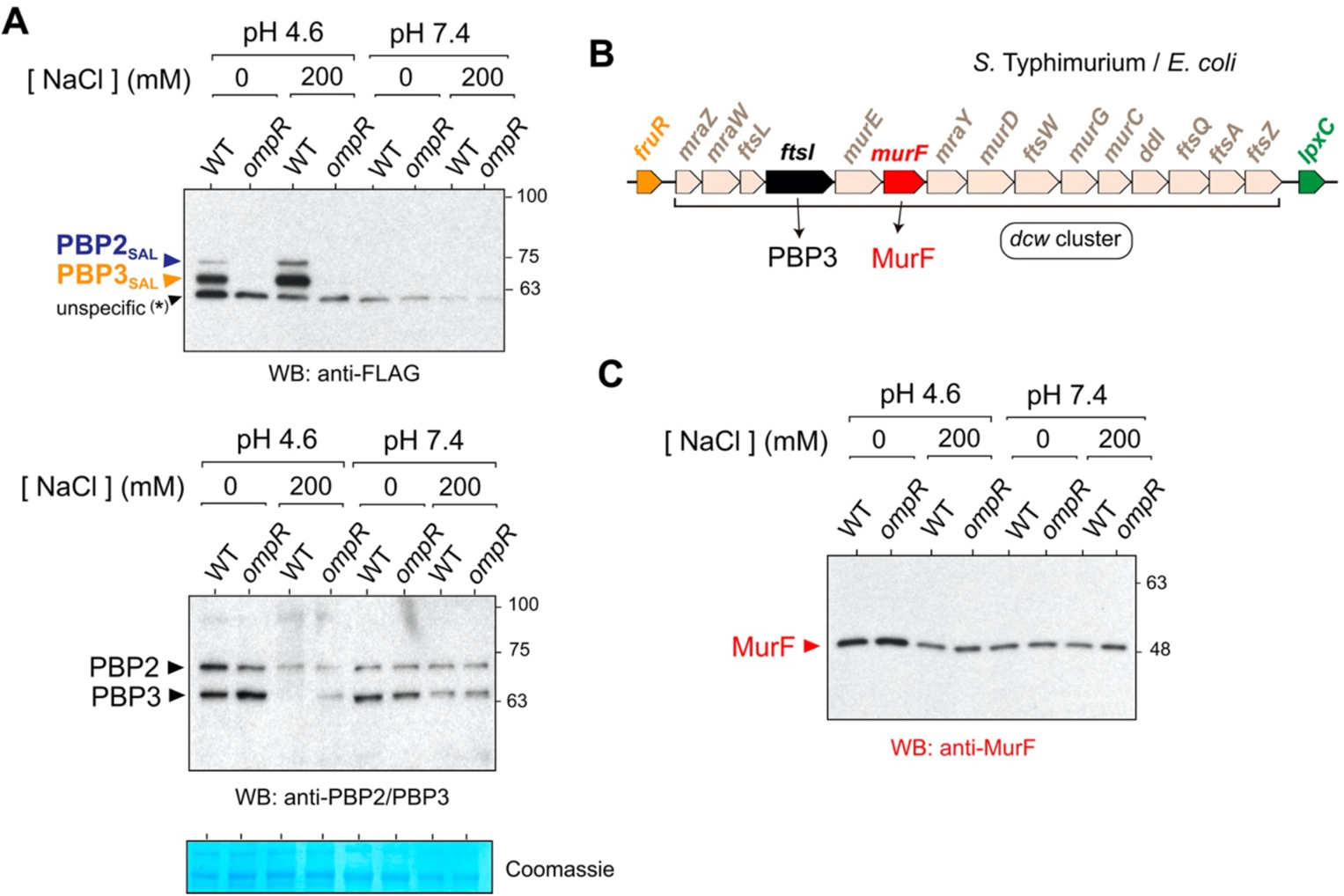
Acid pH and high osmolarity signal *S.* Typhimurium for the precise loss of PBP3. (A) Levels of the enzymatic pairs PBP2SAL/PBP3SAL and PBP2/PBP3 detected in total protein extracts of *S.* Typhimurium wild type and *ompR* strains grown in PCN minimal medium at distinct pH (4.6 and 7.4) and osmolarity (0, 200 mM NaCl). A Coomassie staining of the membrane used in the immunoassays is shown as loading control; (B) Detail of the <u>d</u>ivision and <u>c</u>ell <u>w</u>all (*dcw*) gene cluster that is conserved in *E. coli* and *S.* Typhimurium and functions as an multicistronic operon linked at the transcriptional level. The relative positions of the gene *ftsI*, encoding PBP3, and *murF*, encoding a cytosolic enzyme involved in the synthesis of the PG precursor lipid II, are indicated. (C) MurF levels detected in the total protein extracts used for the determination of PBP2SAL/PBP3SAL and PBP2/PBP3 levels as shown in panel (A). Note that, unlike PBP3, the relative levels of MurF do not drop substantially in high osmolarity and acid pH.

We next monitored in these minimal media conditions the relative levels of MurF, an essential cytosolic enzyme involved in synthesis of the PG precursor lipid II and that is encoded by a gene mapping downstream of PBP3-encoding gene *ftsI.* Both *murF* and *ftsI* map in the <u>d</u>ivision and <u>c</u>ell <u>w</u>all (*dcw*) gene cluster that functions as a single polycistronic transcriptional unit (Fig. 3B). Unlike PBP3, MurF levels did not show substantial changes in response to acid pH and/or high osmolarity, and with no effect for the presence/absence of OmpR (Fig. 3C). Collectively, these observations sustain post-transcriptional negative regulation acting specifically on PBP3 when *S.* Typhimurium encounters acid pH and high osmolarity. Likewise, the data show that upregulation of PBP3_SAL_, triggered solely by acid pH, differs from the elimination of PBP3, which requires both acidity and high osmolarity.

OmpR allows *S.* Typhimurium to divide in an acidic environment with high osmolarity. Since PBP3 is required for cell division, we reasoned that its elimination without having PBP3_SAL_ as counterpart could lead to blockage of such essential process in the cell cycle. This phenotype was expected for the *S.* Typhimurium *ompR* mutant, which only shows trace amounts of PBP3 when growing in minimal medium pH 4.6 and 200 mM NaCl (Fig. 3A). We also sought to determine whether *E.coli*, which has an absolute requirement of PBP3 for division, could negatively regulate its production if exposed to acid pH and high osmolarity. Surprisingly, western blot analyses showed that *E. coli* K-12 strain MG1655, as *S.* Typhimurium, loses PBP3 when growing in minimal medium at pH 4.6 and 200 mM NaCl (Fig. 4A). The loss of PBP3 impacted the cultivability of both *S.* Typhimurium *ompR* mutant and *E. coli* MG1655 in acid pH and high osmolarity although both strains displayed similar growth rate in liquid culture compared to wild type bacteria (Fig. 4B). This latter observation suggested that acidity and high osmolarity did not affect increase in cell mass but promoted a selective loss of PBP3. Such conclusion was further supported by microscopy, which showed that the *S.* Typhimurium *ompR* mutant and wild type *E. coli* MG1655 strains do not divide properly when growing in pH 4.6 and 200 mM NaCl (Fig. 4C). Collectively, these results unveiled the key role that OmpR plays in *S.* Typhimurium for colonizing acidic niches with high osmolarity by inducing the expression of an alternative PG enzyme, PBP3_SAL,_ capable of promoting cell division under those conditions (Castanheira et al., 2017).

**Figure 4.**
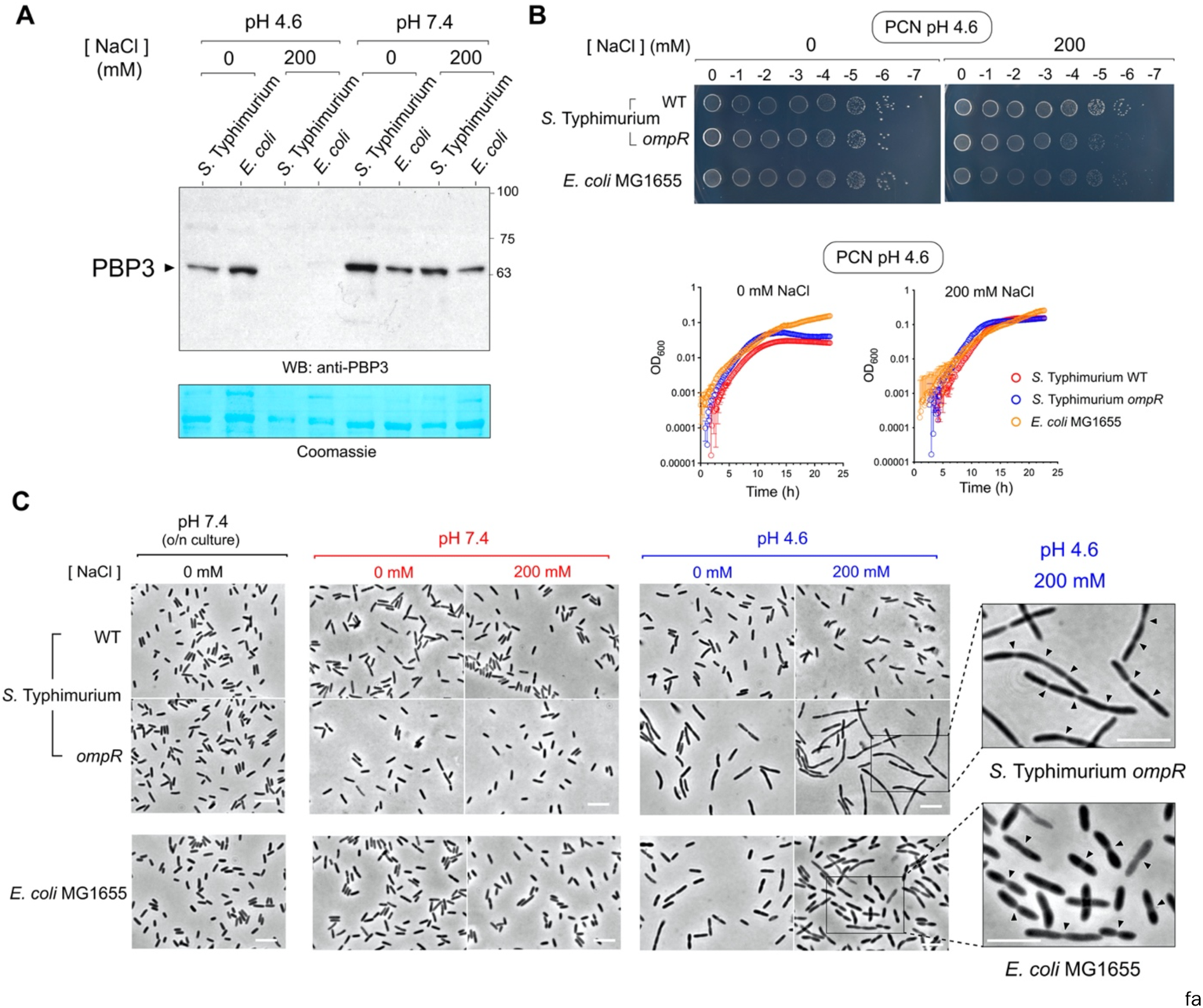
The net production of PBP3 in acid pH and high osmolarity is arrested in both S. Typhimurium and *E. coli*. (A) Comparison of PBP3 levels determined in total protein extracts of *S.* Typhimurium and *E. coli* wild type strains grown in PCN minimal medium at the indicated pH and osmolarity. Note the marked decrease of the enzyme in bacteria exposed to acid pH and high osmolarity; (B) Cultivability of *S.* Typhimurium (wild type and *ompR* strains) and *E. coli* MG1655 wild type on plates of PCN minimal medium at pH 4.6 and the indicated osmolarity (0, 200 mM NaCl). Shown are serial dilutions of an overnight culture obtained in PCN medium 0 mM NaCl pH 7.4. Included are also the growth curves of three strains in liquid PCN medium pH 4.6 with low or high osmolarity (0 or 200 mM NaCl, respectively); (C) Microscopy observation of *S.* Typhimurium (wild type and *ompR* strains) and *E. coli* MG1655 after growing in PCN medium for 4 h at the indicated pH and osmolarity. The inoculum was in all cases obtained in PCN medium pH 7.4, 0 mM NaCl. Shown are also enlargements of *S.* Typhimurium *ompR* and wild type *E. coli* MG1655 cell in acid pH (4.6) and high osmolarity displaying morphological alterations consistent with the loss of PBP3 and, consequently, with the arrest of the cell division process. The arrowheads point to cell division sites in which a small constriction is observed, compatible with the loss of PBP3. Bar, 5 µm. Data shown are representative of a minimum of three independent biological replicates.

The two-component system PhoP-PhoQ does not contribute to the switch of the morphogenetic PG synthases in *S.* Typhimurium. The PhoP-PhoQ two component system is an important regulator modulating the adaptation of *S.* Typhimurium to the intra-phagosomal lifestyle in response to signals like acid pH (Martin-Orozco et al., 2006) and micromolar levels of the divalent cation magnesium (García Véscovi et al., 1996). In extracellular conditions, the *S.* Typhimurium *phoP* mutant showed a slight decrease in the levels of PBP2_SAL_ when growing in acidified LB medium (pH 4.6) and in those of both enzymes, PBP2_SAL_ and PBP3_SAL,_ when infecting fibroblasts (Fig. 1A, 1D).

Based on these data, we monitored the levels of PBP2/PBP3 and PBP2_SAL_/ PBP3_SAL_ in *S.* Typhimurium wild type and *phoP* isogenic strains grown in defined minimal media differing in parameters such as: i) pH (acid 4.6, neutral 7.4); ii) osmolarity (0, 200 mM NaCl); or, iii) magnesium concentrations (8 µM, 10 mM). None of these variables showed a PhoP-dependent regulatory mechanism on any of the four PBPs examined (Fig. 5A-B). Interestingly, the levels of PBP2 and PBP3 dropped in acid pH (4.6) at low magnesium concentration (8 µM) (Fig. 5B), but this change took place independently of PhoP. Taken together, these data ruled out a contribution of the PhoP-PhoQ system in the switch of essential morphogenetic PG synthases occurring when *S.* Typhimurium is exposed to an acidified medium with high osmolarity.

**Figure 5.**
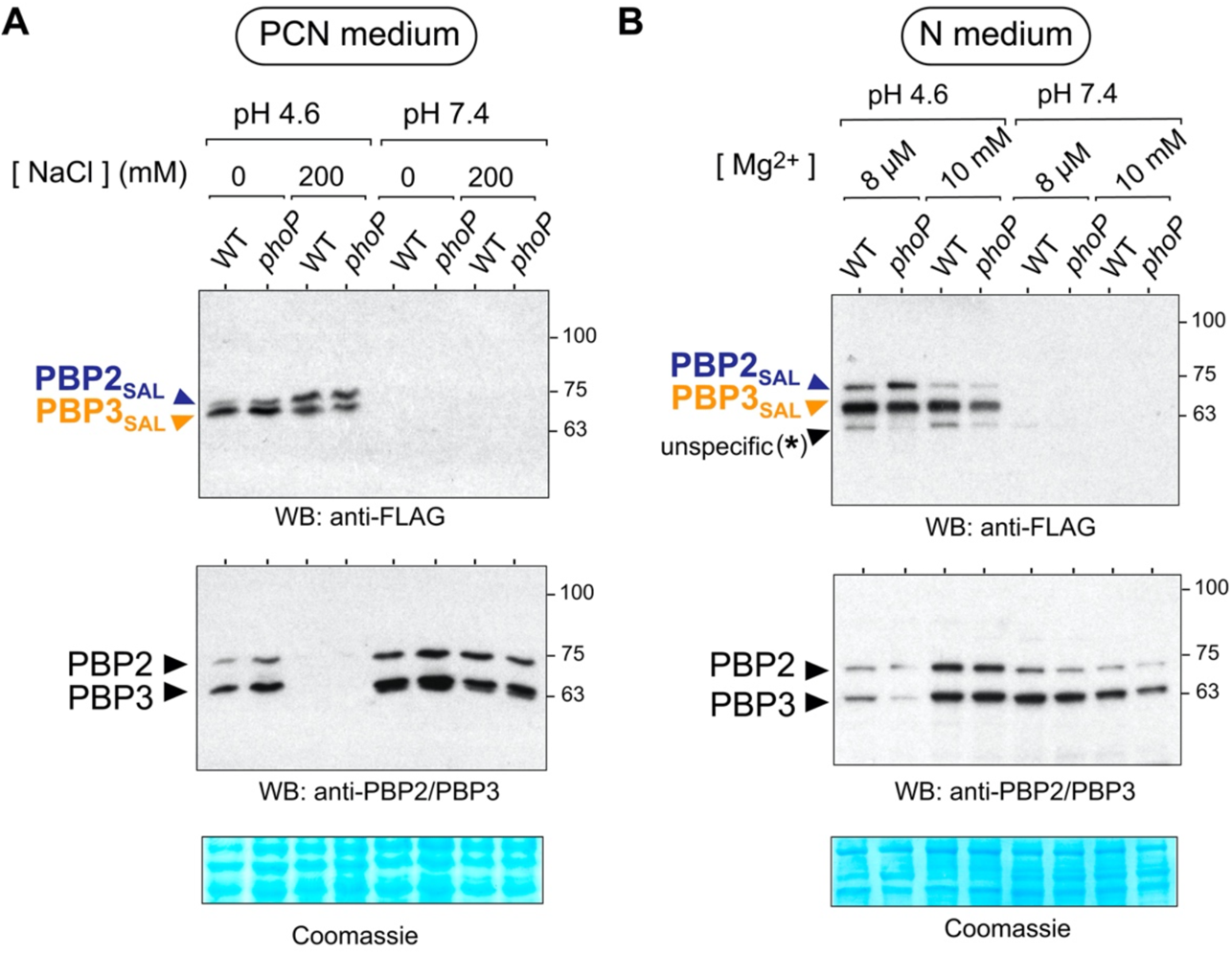
Acid pH and limited amount of magnesium signal *S.* Typhimurium to replace PBP2/PBP3 by PBP2SAL/PBP3SAL in a PhoP-independent manner. (A) Levels of the pair of enzymes PBP2/PBP3 and PBP2SAL/PBP3SAL present in total protein extracts of *S.* Typhimurium wild type and phoP isogenic strains grown in PCN minimal medium at the indicated pH and osmolarity; (B) same as in (A) but in bacteria grown in N minimal media at two different pH (4.6 and 7.4) and magnesium concentrations (8 µM and 10 mM). Note that the condition with low magnesium concentration and acid pH triggers the loss of PBP2 and PBP3 while the production of PBP2SAL and PBP3SAL increases notoriously. Data shown are representative of a minimum of three independent biological replicates.

The protease Prc targets PBP3 and PBP3_SAL_ in the morphogenetic switch and in a pH-dependent manner. In *E. coli*, the protease Prc processes the C-terminal region of PBP3 (Hara et al., 1989; Nagasawa et al., 1989) having as additional substrates other enzymes related to PG metabolism such as the endopeptidase MepS (Spr) (Singh et al., 2015) and the murein transglycosylase MltG (Hsu et al., 2020). The cleavage site recognized by Prc in the PBP3 of *E.coli* (588 residues) is valine-577 (V577) (Nagasawa et al., 1989). Interestingly, this V577 residue is conserved in the PBP3 of S. Typhimurium, and the orthologue PBP3_SAL_, of 581 aa, has also a valine located 11 residues apart of the C-terminus, concretely V570 (Castanheira et al., 2017). To test whether Prc targets any of these two PBPs in *S.* Typhimurium, we determined levels of the precursor and the processed forms of PBP3 and PBP3_SAL_ in wild type and Δ*prc* strains, including also in parallel wild type *E. coli* as additional control.

In acid pH (4.6) and no salt added (0 mM NaCl), the absence of Prc resulted in *S.* Typhimurium in decreased levels of PBP3 (Fig. 6A) whereas in *E. coli* it was clear the presence of two bands, the precursor and mature forms of PBP3 (Fig. 6A). Interestingly, PBP3 is not processed by S. Typhimurium under these conditions (pH 4.6, 0 mM NaCl) (Fig. 6A). As expected, no PBP3 was visualized in any of the strains tested at pH 4.6, 200 mM (Fig. 6A). At neutral pH 7.4, both PBP3 forms (precursor, mature) were visible in S. Typhimurium wild type cells whereas only the full length PBP3 was detected in the Δ*prc* mutant (Fig. 6B). This result unequivocally showed that Prc of *S.* Typhimurium also cleaves PBP3, but only at neutral pH (compare Fig. 6A and 6B). The capacity of *S.* Typhimurium Prc to cleave PBP3 in neutral pH of 7.4 was OmpR-independent regardless of the osmolarity used (Fig. 6C).

**Figure 6.**
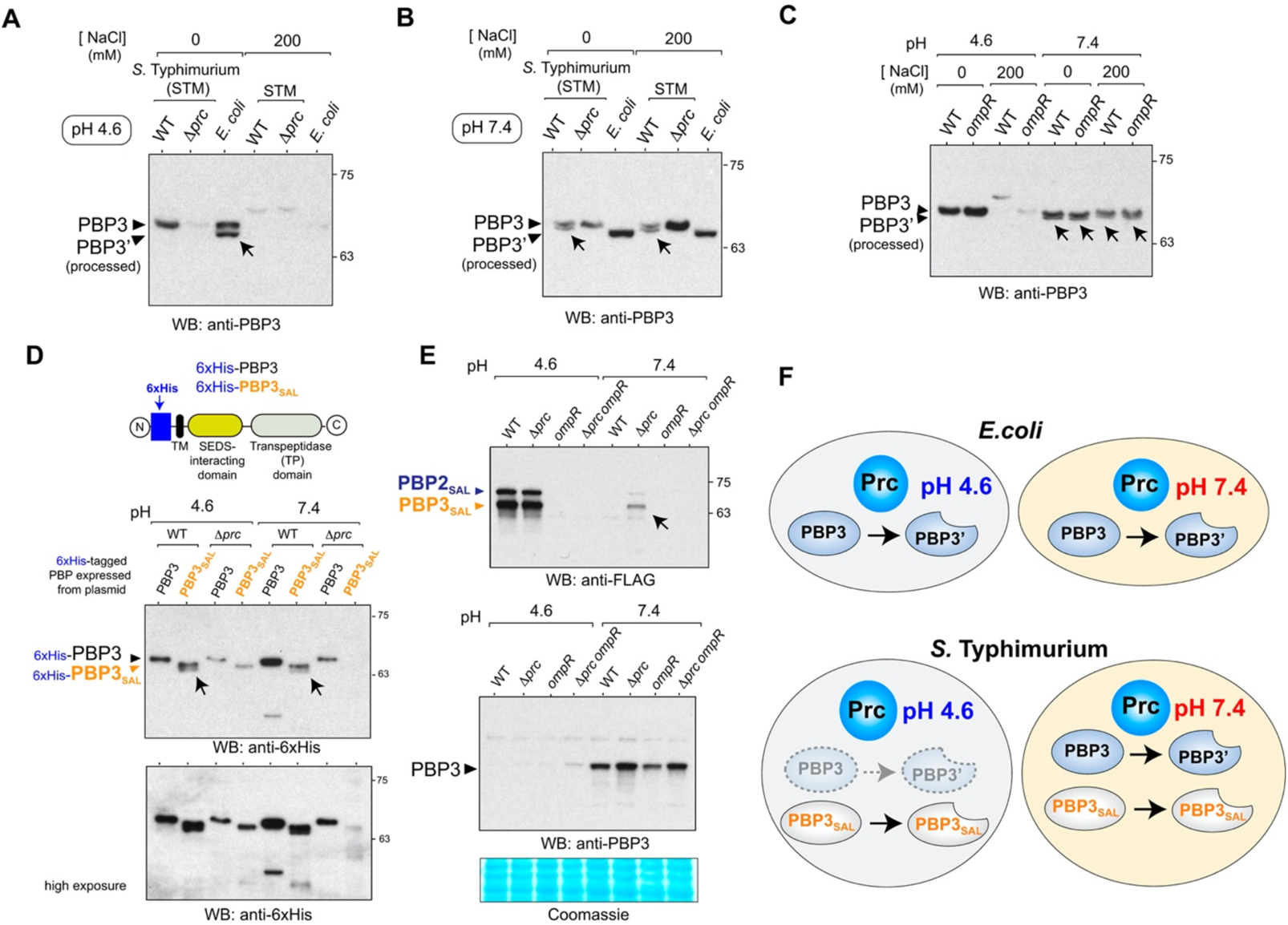
*S.* Typhimurium exploits the protease Prc to target selectively PBP3 or PBP3SAL at distinct pH. (A) Processing of PBP3 is not observed in wild type *S.* Typhimurium at acid pH (4.6) whereas this PBP3 is cleaved by Prc in *E. coli* in the same condition (PCN medium, pH 4.6, 0 mM NaCl). Note the loss of PBP3 concomitant to the increase in osmolarity (200 mM NaCl) and the loss of PBP3 in the *S.* Typhimurium Δ*prc* mutant at low osmolarity. Arrow indicates the mature processed form of PBP3; (B) S. Typhimurium Prc cleaves PBP3 at neutral pH (7.4) irrespective of the amount of salt added to the medium (0, 200 mM NaCl). The PBP3 form observed for *E. coli* in these assays is processed as inferred from its higher electrophoretic mobility. Arrows point to the processed mature form of PBP3; (C) The protease activity of *S.* Typhimurium Prc over PBP3 at neutral pH (7.4) occurs independently of OmpR and osmolarity. Arrows point to the processed mature form of PBP3; (D) Prc cleaves a 6xHis variant of PBP3SAL tagged at the N-terminus expressed from plasmid at both, acid (4.6) and neutral (7.4) pH. Arrows point to the processed form of PBP3SAL. Unlike the case of 6xHis-PBP3SAL and the PBP3 expressed from its native chromosomal location [see panel (B)], the 6xHis-PBP3 tagged variant expressed from plasmid is not cleaved by Prc; (E) Detection of minor amounts of PBP2SAL/PBP3SAL at neutral pH in the absence of Prc (arrow), may indicate the existence a quality control exerted by this protease at this pH; (F) Diagram depicting the preferred recognition of *S.* Typhimurium for PBP3SAL at acid pH and for PBP3 at neutral pH although also showing the capacity of the protease to target PBP3SAL if produced at neutral pH.

With regards to PBP3_SAL_ and considering the presence of a residue V570 putatively recognized by Prc, we assessed whether it could be cleaved by this protease at the C-terminus. To this aim, we used PBP3_SAL_ and PBP3 versions bearing a 6xHis tag at the N-terminus (Castanheira et al., 2020). These two variants, 6xHis-PBP3 and 6xHis-PBP3_SAL,_ were expressed from plasmid in wild type and Δ*prc* strains at acid (pH 4.6) and neutral (7.4) pH (Fig. 6D). These assays showed that Prc cleaves 6xHis-PBP3_SAL_ at the C-terminus at both pH whereas no processing was observed for 6xHis-PBP3 (Fig. 6D). This result in PBP3 suggests that the modification at the N-terminus of this enzyme, which faces the cytosol, could alter its recognition in the periplasm by the protease, an effect not seen in unmodified PBP3 (Fig. 6B).

Given the capacity of Prc to cleave the C-terminus of PBP3_SAL_ (Fig. 6D), we assessed whether this protease could play a role in *S.* Typhimurium in preventing the presence of this enzyme when the pathogen grows in neutral pH. The lack of Prc resulted in the detection of a small amount of PBP3_SAL_ at pH 7.4 (Fig. 6E). The detection of these low amounts of PBP3_SAL_ at neutral pH in the absence of the protease was, however, OmpR-dependent (Fig. 6E). Taken together, these data demonstrated that PBP3 and PBP3_SAL_ are substrates of Prc in *S.* Typhimurium although with a clear dependence on the pH. Thus, whereas Prc cleaves PBP3 only at neutral pH, it processes PBP3_SAL_ in both the physiological condition in which its production is triggered by OmpR (acid pH of 4.6) and in neutral pH. Unlike Prc of *S.* Typhimurium, this protease in *E. coli* cleaves PBP3 at both acid and neutral pH (Fig. 6F).

## Discussion

Morphogenesis has been mostly investigated in bacteria exhibiting rod or coccoid shapes in defined (controlled) environmental conditions (Daitch and Goley, 2020; Egan et al., 2020; Radkov et al., 2018) and in organisms exhibiting morphological plasticity (Caccamo and Brun, 2018). These latter include those generating differentiated cells, subcellular structures like stalks and branching or curved shapes (Caccamo and Brun, 2018; Khanna et al., 2020). The PG determines cell shape and some studies demonstrate that specific enzymes acting on this macromolecule are responsible for morphological plasticity (Caccamo and Brun, 2018). Inversely, *S. enterica* serovar Typhimurium represents what to our knowledge is the only case known to date involving two morphogenetic machineries with distinct PG synthases (PBP2/PBP3 and PBP2_SAL_/PBP3_SAL_) to generate the same (rod) cell shape.

To get insights into how *S.* Typhimurium regulates this switch of morphogenetic machineries, we initially inspected expression levels of the PBP2_SAL_- and PBP3_SAL_-encoding genes *SL1344_1845* and *SL1344_1765* in transcriptomic datasets. Data obtained from bacteria infecting cultured macrophages (Eriksson et al., 2003), fibroblasts (Núñez-Hernández et al., 2013), epithelial cells (Hautefort et al., 2008) or exposed to varied stress conditions in laboratory media (Kroger et al., 2013), indicated that these two genes are poorly transcribed in all conditions tested. Only in a few cases, increased transcription, in the order of 3-fold, was registered for the PBP3_SAL_-encoding gene in bacteria recovered from macrophages or exposed to high salt (0.3 M NaCl, 10 min), acid (pH 5.8) or bile (3% bile, 10 min) (Kroger et al., 2013). This reduced transcriptional activity contrasts with the relatively high amounts of PBP2_SAL_ and PBP3_SAL_ that are detected by western blot [(Castanheira et al., 2020, 2017), this study].

Our data unequivocally show that OmpR controls the production of these two PG enzymes and that acid pH is necessary and sufficient for such positive regulation. OmpR, which was first identified as a regulator of the levels of the two major outer membrane proteins, OmpC and OmpF (Taylor et al., 1981), was latter shown to control these porins via regulatory small RNAs (sRNAs) like MicF and MicC that target the *ompF* and *ompC* transcripts, respectively (Vogel and Papenfort, 2006). OmpR induces transcription of two other sRNAs, OmrA and OmrB, which control post-transcriptionally the stability of multiple mRNA, including the *ompR-envZ* transcript as a feedback auto-regulatory loop (Brosse et al., 2016). Therefore, a recurrent theme in the mechanisms that OmpR uses as regulator is the simultaneous action at transcriptional and post-transcriptional levels by controlling transcription of the target mRNA and sRNA that will regulate post-transcriptionally such target (Brosse et al., 2016). In this scenario, it is tempting to speculate on sRNAs modulating the stability and/or translation of the mRNAs encoding PBP2_SAL_ and PBP3_SAL,_ a possibility that merits to be investigated in future work.

When compared to other virulence-related regulators, it is clear that OmpR plays a master role in controlling the production of PBP2_SAL_ and PBP3_SAL,_ both in laboratory media and inside host cells. This regulation could be coordinated with that PhoP may exert on PBP2_SAL_ in extracellular bacteria (Fig. 1A) or in intracellular bacteria for the case of PBP2_SAL_ and PBP3_SAL_ (Fig. 1D). OmpR and PhoP control expression of horizontally acquired foreign genes and crosstalk in regulating SPI-2, the key pathogenicity island exploited by *S.* Typhimurium for a successful intracellular infection (Fass and Groisman, 2009; Kim and Falkow, 2004; Lee et al., 2000; Liew et al., 2019; Worley et al., 2000). Our results, however, show that SsrB, absolutely required for expression of SPI-2 genes, plays no role in the production of PBP2_SAL_ and PBP3_SAL_. This finding suggests that PBP2_SAL_/PBP3_SAL_ and SPI-2 might be independent targets in the OmpR and PhoP regulons. The data obtained with the transcriptional reporter fusions showed that OmpR acts canonically on the *SL1344_1845* and *SL1344_1765* promoters recognizing sequences that match the consensus TGTWACAW (Perkins et al., 2013). The OmpR box present in the promoter of the PBP3_SAL_-encoding gene has a mismatch respect the consensus, G in fifth position (TGTTGCAA), which may explain why its reporter was expressed at lower levels than that of PBP2_SAL_ (Fig. 2). Importantly, the expression of these reporters was independent of PhoP. Based on these findings, the positive regulation that PhoP may exert on PBP2_SAL_ and PBP3_SAL_ could be indirect, e.g. via regulatory sRNAs targeting the respective mRNAs. Another relevant finding in these assays was the reproducibility of the reporter expression pattern in a heterologous system of *E. coli,* showing the same dependence on acid pH as in *S.* Typhimurium. Based on these findings, we therefore conclude that OmpR binding to the PBP2_SAL_ and PBP3_SAL_ promoters is pH-sensitive.

Acid pH and high osmolarity are signals perceived by the sensor EnvZ to activate OmpR (Chakraborty and Kenney, 2018; Kenney, 2019; Quinn et al., 2014). Genome-wide expression analyses show that each signal by separate triggers only partly-overlapping OmpR regulons (Chakraborty and Kenney, 2018). Our analyses of PBP2_SAL_ and PBP3_SAL_ at the protein level supports this view since acid pH of 4.6 as single signal is sufficient to detect high levels of the two enzymes. Low magnesium levels in the order of micromolar, known to signal the sensor PhoQ to activate PhoP, are however not required for PBP2_SAL_ and PBP3_SAL_ production (Fig. 5B). These data are consistent with a regulation of PBP2 _SAL_ and PBP3_SAL_ triggered essentially by acidification of the cytoplasm mediated by the transcriptional regulator OmpR (Kenney, 2019). Cytoplasm acidification is known to stimulate the cytoplasmic domain of the sensor PhoQ leading to increased expression of PhoP-activated genes (Choi and Groisman, 2016), some of which could be potentially linked to ensure the required levels of PBP2_SAL_ and PBP3_SAL_, especially in response to intracellular cues (Fig. 1D).

We were also much interested in getting insights into how *S.* Typhimurium diminishes PBP2 and PBP3 levels in infection conditions given these are essential enzymes in *E. coli*. The elimination of PBP2 and PBP3 was probably crucial in evolutionary terms by acting as selective force that facilitated the fixation of a second morphogenetic machinery, fully active in acid pH and high osmolarity. To recall that in the laboratory the complete replacement of PBP2/PBP3 by PBP2_SAL_/PBP3_SAL_ that is observed *in vivo* in mouse tissues (Castanheira et al., 2020) is only partially reproduced when using the acidified PCN medium. This is more notorious for PBP2, for which a small amount of the enzyme is still detected in acid pH despite the *de novo* synthesis of PBP2_SAL_. Fortunately, that was not the case for PBP3/PBP3_SAL_, in which the acidified PCN medium with high amount of salt (200 mM NaCl) led to a complete interchange of both enzymes in *S.* Typhimurium. We are currently taking caution about this coexistence of morphogenetic enzymes since in certain media, as an acidified nutrient rich LB medium, both sets of enzymes (PBP2/PBP2_SAL_ and PBP3/PBP3_SAL_) are detected at similar levels (Castanheira et al., 2017), a situation that might not occur in natural habitats colonized by this pathogen.

The data presented here also implicate OmpR in the elimination of PBP2 and PBP3 in response to acid pH although, in this case, additional regulatory factors like high osmolarity, the periplasmic protease Prc and a low concentration of Mg^2+^ were equally involved (Fig. 7). Surprisingly, this intricate “negative” regulatory network is conserved in *E. coli*, at least for PBP3, as this organism loses this enzyme and is unable to divide if exposed to acid pH (4.6) and 200 mM NaCl. Intriguingly, uropathogenic *E. coli* (UPEC) undergoes filamentation during urinary tract infections as a strategy to subvert host immune defences (Horvath et al., 2011; Justice et al., 2008, 2006). This rod-to-filament morphological transition is stimulated during the intracellular stage, in which probably bacteria are exposed to an acid environment. A recent study in *E. coli* has also provided evidence of an sRNA, FtsO, mapping internally to the PBP3-encoding gene *ftsI* (Adams et al., 2021). Despite not knowing how FtsO is generated, it modulates the levels of the *ompC* transcript via the sRNA RybB, and its presence supports the existence of a post-transcriptional regulatory mechanism that could act specifically to regulate PBP3 levels concomitantly to those of the outer membrane protein OmpC. The loss of PBP3, detectable also in *E. coli*, underscores how relevant was for *S.* Typhimurium the acquisition of PBP3_SAL_ to colonize eukaryotic acidic phagosomes ensuring the maintenance of the progeny.

**Figure 7.**
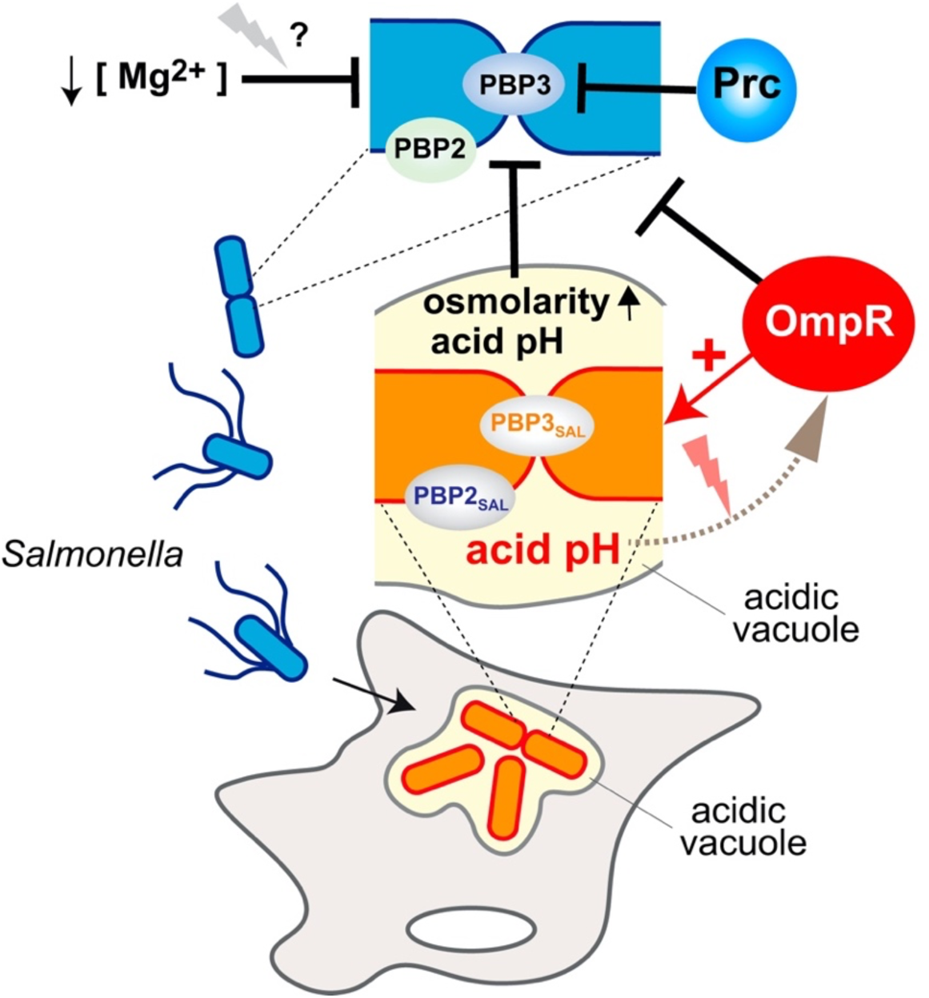
Hypothetical model integrating the regulatory factors that promote in *S.* Typhimurium the switch of PBP2/PBP3 by PBP2SAL/PBP3SAL during the transition to the intracellular lifestyle inside acidic vacuoles of the eukaryotic cell. Key regulatory events include the absolute requirement of OmpR for the production of PBP2SAL/PBP3SAL by intracellular bacteria in response to acid pH and the multifactorial nature that characterizes the negative regulation over PBP2/PBP3 in which OmpR, acid pH, high osmolarity, the protease Prc and a low magnesium concentration may contribute synergistically.

Our study has also shed light into the activity of the protease Prc on PBP3 and PBP3_SAL._ Our electrophoretic analyses convincingly showed that Prc can cleave both substrates in *S.* Typhimurium. The significance of the cleavage exerted by the Prc protease in the activity of PBP3 (and PBP3_SAL_) remains however unclear. Thus, although displaying temperature sensitive and outer membrane leaky phenotypes, *E. coli* mutants lacking Prc are viable [reviewed in (Hara et al., 1996)]. *S.* Typhimurium *prc* null mutants are also viable (Hernández et al., 2013), indicating that processing of PBP3 and PBP3_SAL_ is not absolutely essential for cell division. Nonetheless, the data obtained in *S.* Typhimurium point to a putative relevant role of Prc in the replacement of the morphogenetic machineries. Unlike its *E. coli* counterpart, Prc of *S.* Typhimurium does not cleave PBP3 in acid pH whereas it cleaves this substrate at neutral pH. This behaviour contrasts with its other substrate, PBP3_SAL_, which is cleaved at both acid neutral pH when expressed from plasmid. We also noted that, only for the case of PBP3, the addition of a 6xHis tag in the N-terminus impaired cleavage of this specific enzyme at neutral pH (Fig. 6D), a result that is not easy not be interpreted given the lack of studies addressing the role of the short N-terminal region of the enzyme that faces the cytosol. Tagging this region might therefore has consequences in how PBP3 is recognized by the protease in the periplasm, an effect that may not occur for PBP3_SAL_.

An unexpected data was the appearance of small amounts of PBP3 and PBP3_SAL_ in their “non-natural” conditions, PBP3_SAL_ in neutral pH and PBP3 in acid pH and high osmolarity conditions, when Prc was lacking (Fig. 6E). This result might indicate that Prc exerts a quality control aborting two phenomena: i) the escape that may exist in the negative regulation that OmpR imposes over PBP3 in acid pH; and, ii) the fortuitous production of PBP2_SAL_ and PBP3_SAL_ at neutral pH dependent on OmpR (Fig. 6E). Whether Prc exerts this role directly on these PG enzymes, is at present unknown.

Our study therefore unravels the first regulatory rules controlling the switch of two morphogenetic machineries involving essential PG synthases that takes place when *S.* Typhimurium senses acid pH and high osmolarity. Given that adaptation to these conditions is a requisite for successful infection of eukaryotic cells, it is tempting to speculate that this exchange of PG enzymes might be interconnected to virulence functions activated by *S.* Typhimurium in response to intra-phagosomal cues.

## Materials and Methods

### Bacterial strains, growth media

Bacterial strains and plasmids used are listed in Table S1. All *S. enterica* serovar Typhimurium (*S.* Typhimurium) strains used are derivates of virulent wild-type strain SV5015, an His+ prototroph derivate of virulent strain SL1344 (Vivero et al., 2008). Strains were grown in Luria-Bertani (LB) broth for conditions with high amount of nutrients. LB broth is composed of 1% (w/v) casein peptone, 0.5% (w/v) yeast extract and 0.5% (w/v) sodium chloride. For nutrient-limiting conditions simulating the intra-phagosomal environment, strains were grown in phosphate-carbon-nitrogen (PCN) minimal medium (Deiwick and Hensel, 1999) or N minimal medium (Nelson and Kennedy, 1971). The composition of PCN medium is: 4 mM Tricine [N-[Tris (hydroxymethyl) methyl]glycine], 0.1 mM FeCl_3_, 376 µM K_2_SO_4_, 15 mM NH_4_Cl, 1 mM MgSO_4_, 10 µM CaCl_2_, 0.4% (w/v) glucose, 0.4 mM K_2_HPO_4_/KH_2_PO_4_, and micronutrients (Deiwick and Hensel, 1999). 200 mM NaCl was added to PCN to test response to high osmolarity. N minimal medium is composed of 5 mM KCl, 7.5 mM (NH_4_)_2_S0_4_, 0.5 mM K_2_S0_4_, and 1 mM KH_2_PO_4_ supplemented with 0.4% (w/vol) glucose as carbon source, 0.1% (w/vol) casaminoacids and 10 mM or 8 µM MgCl_2_ (high or low Mg^2+^ concentrations, respectively) (Snavely et al., 1991). When necessary, pH was buffered with 80 mM MES [2-(N-morpholino) ethanesulfonic acid] adjusted to the desired value with NaOH. To grow strains bearing genetic elements conferring antibiotic-resistance, media were supplemented with chloramphenicol (10 µg/mL), kanamycin (30 µg/mL), or ampicillin (100 µg/mL). For induction of gene expression from the P*lac* promoter in pAC-His plasmids (see Table S1), 10 µM IPTG (Isopropyl β-D-1-thiogalactopyranoside) was added. Plasmids expressing 6xHis-PBP3 and 6xHis-PBP3sal variants tagged in the N-terminus has been previously described (Castanheira et al., 2020).

### Construction of chromosomal epitope-tagged genes

Strains carrying chromosomal 3×FLAG-epitope-tagged genes were constructed using the method described by Uzzau et al. (Uzzau et al., 2001). The phage P22 HT 105/1 *int*201 (Schmieger, 1972) was used for transductional crosses to mobilize mutant alleles for strain construction. The P22 HT transduction protocol was described elsewhere (Garzón et al., 1995). To obtain phage-free isolates, transductants were purified by streaking on green plates, prepared according to Chan et al. (Chan et al., 1972).

### Construction of a *S.* Typhimurium Δ*prc* mutant expressing PBP2_SAL_-3xFLAG and PBP3_SAL_-3xFLAG tagged variants

A P22 HT 105/1 *int*201 lysate was first obtained from *S.* Typhimurium strain SV6246 (Δ*prc*::Km^R^) (Hernández et al., 2013), a gift from J. Casadesús (University of Seville, Spain). This deletion mutant harbors a null allele lacking the entire *prc* coding sequence (Hernández et al., 2013). The phage lysate was used to transduce strain MD5064 (PBP2_SAL_-3xFLAG PBP2_SAL_-3xFLAG) to obtain strain MD5416 (Δ*prc*::Km^R^ PBP2_SAL_-3xFLAG PBP2_SAL_-3xFLAG) (Table S1).

### Growth conditions for preparation of total protein extracts

When using PCN minimal medium, *S.* Typhimurium and *E. coli* strains were first grown overnight with agitation at 37°C, pH 7.4 without NaCl supplementation. To monitor the effect of osmolarity on the production of PBPs, the bacteria from overnight cultures were washed [4,300 x *g*, 2 min, room temperature (RT)] with PCN medium at the conditions of interest: pH 4.6 or 7.4, and NaCl concentrations of 0 mM or 200 mM. The initial optical density at 600 nm (OD_600_) of the culture was established at 0.02 in 20 mL cultures. When using N minimal medium, *S.* Typhimurium strains were grown overnight with agitation at 37°C, pH 7.4 with a MgCl_2_ concentration of 10 mM. Overnight cultures were then washed (4,300 x *g*, 2 min, RT) with N minimal medium pH 4.6 or 7.4, and MgCl_2_ concentration of 8 µM or 10 mM, according to culture growth conditions, and diluted to an initial OD_600_ of 0.02 in 20 mL cultures. In minimal media, bacteria were collected after 4 h of growth at 37°C in agitation (150 rpm). When using LB, bacteria were first grown overnight in agitation at 37°C in non-buffered neutral pH LB. Overnight cultures were diluted to an initial OD_600_ of 0.02 in 20 mL cultures with LB or LB pH 4.6 buffered with MES and bacteria collected at 3 h post-inoculation. To trigger production of PBP3 or PBP3_SAL_ from pAC-His plasmids (see Table S1), strains were grown overnight with agitation (150 rpm) at 37°C in LB and diluted to an OD_600_ of 0.02 in fresh LB or LB pH 4.6 buffered with MES. Strains were then grown at 37°C for 1.5 h, induced with 10 µM IPTG for gene expression and grown for further 1.5 h before being collected for total protein extraction. In all growth conditions, bacteria were harvested by centrifugation at 18,000 × *g*, 5 min, 4°C, washed twice in phosphate-buffered saline (PBS) and resuspended in 100 µL of Laemmli lysis buffer per optical density unit.

### Large-scale infection of cultured fibroblasts

Large-scale infections were performed to obtain protein from intracellular bacteria, essentially as previously described (Núñez-Hernández et al., 2013). NRK-49F normal rat kidney fibroblasts (ATCC CRL-1570) were seeded in Nunc Square BioAssay Dishes (Thermo Scientific, ref. 166508) with 50 mL Dulbecco’s Modified Eagle Medium (DMEM) culture medium supplemented with 5% (v/v) foetal bovine serum (FBS) and 4 mM L-glutamine to a confluence of 80% (~5.0 x 10^7^ cells). The fibroblast culture was infected with the different *S.* Typhimurium strains, previously grown overnight at 37°C in static LB culture, for 40 min at an MOI of 10:1 (bacteria: fibroblast). At this time post-infection, cells were washed two times with prewarmed complete PBS solution (PBS pH 7.4 with 0.9 mM CaCl_2_ 0.5 mM MgCl_2_) and then incubated in fresh DMEM-5% FBS culture medium containing 100 µg/mL of gentamicin until 2 h post-infection. The culture medium was then replaced with fresh DMEM-5% FBS medium containing 10 µg/mL gentamicin until 8 h post-infection. At that time, the infected fibroblasts were washed five times with 20 mL cold complete PBS and lysed in 17 mL of a solution containing 1 % (v/v) pH 6.6-7.9 basic phenol, 19 % (v/v) ethanol and 0,4 % (w/v) SDS in water. A volume of 1.2 µL DNase (10 mg/mL) was added to each plate. After 30 min of incubation at 4°C, the lysate was collected in 40 mL polypropylene tubes (Sorvall) and centrifuged (27,500 x *g*, 4°C, 30 min). The resultant pellet was washed twice with 1 mL of a 1% basic phenol, 19% ethanol solution (29400 x *g*, 15 min, 4°C). Intracellular bacteria were resuspended in 40 µL Laemmli lysis buffer.

### Immunoblot assays

For SDS/PAGE, samples were incubated at 100°C for 5-10 min and centrifuged to remove cell debris (6,800 x *g*, 5 min, RT) for SDS/PAGE. Proteins resolved by gels were transferred to polyvinylidene difluoride (PVDF) membranes and incubated with the corresponding antibodies. Detection was performed by Clarity Western ECL Substrate chemioluminescence kit (Bio Rad, ref. 1705061). Once used for the Western blot, the PVDF membranes were stained with Coomassie solution to confirm proper adjustment of samples except samples from intracellular bacteria, in which immunodetection of DnaK chaperone was made. Proteins were routinely resolved in 6% polyacrylamide gels. To increase the separation of mature and processed PBP3 forms, 11% gels for the Tricine-electrophoresis system (Schägger and von Jagow, 1987), were used. Similarly, to increase separation between PBP2_SAL_ and PBP3 _SAL_, 4-20% Mini-PROTEAN TGX Precast Protein gels (BioRad, ref. 4561096), were used. The following antibodies were used as primary antibodies: mouse monoclonal anti-Flag (M2 clone, 1:5000; Sigma), mouse monoclonal anti-6xHis (1:2500; R&D Systems), rabbit polyclonal anti-PBP2 (1:1000; lab collection) (Castanheira et al., 2020), rabbit polyclonal anti-PBP3 (1:1000; lab collection) (Castanheira et al., 2017) and, mouse monoclonal anti-DnaK (clone 8E2/2, 1:10000; Enzo Life Sciences). Goat polyclonal anti-mouse (ref. 1706516, 1:20000, BioRad) and anti-rabbit IgG (ref. 1706515, 1:30000, BioRad) conjugated to horseradish peroxidase (Bio-Rad), were used as secondary antibodies.

### Phase-contrast microscopy

Bacteria in overnight cultures were centrifuged (4,300 × *g*, 2 min, RT), washed in PCN medium with the corresponding pH (4.6 or 7.4) and salt concentration (0 mM or 200 mM NaCl) and used to inoculate the respective media to an initial optical OD_600_ of 0.02. After 4 h growing with agitation (150 rpm) at 37°C, bacteria were harvested (6,800 × *g*, 4 min, 4°C), washed twice in PBS buffer, fixed with 3% paraformaldehyde (PFA) for 10 min and adjusted to a final paraformaldehyde (PFA) concentration of 1%. For microscopy, fixed bacteria were centrifuged (4,300 × *g*, 4 min, RT) and resuspended in an equal volume of PBS. A volume of 30 µL was dropped on poly-L-Lys pretreated coverslips and incubated for 10 min at RT. Attached bacteria were washed four times with PBS and the coverslip mounted on slides using ProLong Gold Antifade (Molecular Probes). Images were acquired on an inverted Leica DMI 6000B microscope with an automated CTR/7000 HS controller (Leica Microsystems) and an Orca-R2 charge-coupled-device (CCD) camera (Hamamatsu Photonics).

### Construction of reporter plasmids involving PBP2_SAL_ and PBP3_SAL_ promoter regions

To monitor activity of the predicted promoter regions for the genes encoding PBP2_SAL_ and PBP3_SAL_ a vector with the backbone of plasmid pGEN222 (Galen et al., 1999) bearing an ovoalbumin (OVA)-encoding gene under the P*ompC* promoter and a gene expressing GFP^TCD^ variant (Corcoran et al., 2010), was used. This vector, kindly provided by L.A. Fernández (CNB-CSIC, Madrid, Spain), is named pGEN222-*ova-gfp*. Regions encompassing 250 pb upstream the start codon of PBP2_SAL_- and PBP3_SAL_-encoding genes (*SL1344_1845* and *SL1344_1765*, respectively) were amplified from *S.* Typhimurium SV5015 genomic DNA (gDNA) using the primers Ppbp2sal_EcoRI_NotI_Fw/Ppbp2sal_EcoRI_NotI_Rv and Ppbp3sal_EcoRI_NotI_Fw/ Ppbp3sal_EcoRI_NotI_Rv (Table S2). The PCR products containing the PBP2_SAL_ and PBP3_SAL_ promoter regions were digested with EcoRI and NotI and ligated in an EcoRI/NotI-digested pGEN222-*ova-gfp* replacing, as a result, the p*ompC-ova* region. As negative control, this p*ompC-ova* region was excised with EcoRI/NotI from pGEN222-*ova-gfp* and the plasmid was religated. To this aim, pGEN222-*ova-gfp* digested with EcoRI and NotI was incubated with T4 DNA polymerase (Roche) for 15 min at 12°C, then 2.5 µL of EDTA-Na^+^ 0.1 M were added. The mixture was re-incubated 20 min at 75°C and the purified DNA product further incubated overnight at 16°C with a T4 DNA ligase (Roche).

### Construction of mutated versions of the OmpR boxes in the PBP2_SAL_ and PBP3_SAL_ promoters

An annealing extension procedure was used for generating PBP2_SAL_ and PBP3_SAL_ promoters with altered OmpR boxes. These regions were amplified in two fragments from plasmids pGEN222(Δ*ompC*-*ova*)-P*PBP2_SAL_*-*gfp*^TCD^ and pGEN222(Δ*ompC*-*ova*)-P*PBP3_SAL_*-*gfp*^TCD^, respectively. Fragments with mutations at putative OmpR-binding sites were amplified using the following primers: (i) pPBP2sal_mut_Fw/pPBP2sal_EcoRI_NotI_Rv and pPBP2sal_mut_Rv/ pPBP2sal_EcoRI_NotI_Fw for the PBP2_SAL_ promoter; and, (ii) pPBP3sal_mut_Fw/pPBP3sal_EcoRI_NotI_Rv and pPBP3sal_mut_Rv/pPBP3sal_EcoRI_NotI_Fw, for the PBP3_SAL_ promoter (Table S2). Then, the fragments for each construction were co-incubated for 5 cycles of annealing extension and the resulting full-length fragment were amplified using external primers Ppbp2sal_EcoRI_NotI_Fw/ Ppbp2sal_EcoRI_NotI_Rv and Ppbp3sal_EcoRI_NotI_Fw/ Ppbp3sal_EcoRI_NotI_Rv for 30 cycles. The resulting fragments were digested with EcoRI and NotI restriction enzymes, purified and ligated in a EcoRI-NotI digested pGEN222-derivative vector. All constructs were sequenced with primer pGEN222_Fw (Table S2) to rule out the presence of undesired mutations. The reporter plasmids vectors harboring the mutated versions of the PBP2_SAL_ and PBP3_SAL_ promoters were electroporated into the desired *S.* Typhimurium and *E. coli* strains.

### Monitoring of promoter activity with reporter plasmids expressing GFP^TCD^

The *S.* Typhimurium and *E. coli* strains harboring the reporter plasmids with the PBP2_SAL_ and PBP3_SAL_ promoter regions were grown overnight in PCN minimal medium pH 7.4, 200 mM NaCl at 37°C, then washed once (4,300 x *g*, 2 min, RT) with PCN 200 mM NaCl and pH of either 4.6 or 7.4 and, finally resuspended in these respective media at an initial OD_600_ of 0.02. These cultures were transferred with volumes of 180 µL per well in triplicate into 96-well flat clear bottom black polystyrene TC-treated microplates (Corning®, ref. 3904). Bacteria were incubated at 37°C for 18 h with 20 sec of orbital agitation every 20 min, followed by OD_600_ and fluorescence measurements in a Spark^®^ microplate multimode reader (TECAN). Fluorescence units (FU) were measured with wavelength excitation at 475/15 nm and emission at 510/15 nm. The background values of both optical density and fluorescence were subtracted in the values of the bacterial samples. Fluorescence units were finally corrected by optical density for graphical representation.

### DNA techniques

All primers used in this study are listed in Table S2. PCR was performed using Q5 polymerase (New England Biolabs) according to manufacture instructions. PCR fragments were purified using the NucleoSpin Gel and PCR Clean-up kit (Macherey-Nagel, ref. 740609.50). Plasmids were purified using the NZYMiniprep kit (NZYTech, ref. MB01001).

### Measurements and statistical analysis

Intensity of protein bands obtained in the immunoassays was measured with Fiji distribution of ImageJ2 (version 1.52i) (Schindelin et al., 2012). Similar band expositions for reference strain were selected to increase the comparability between data of distinct biological replicates. Data were analyzed with GraphPad Prism software v8.0 (GraphPad Inc. San Diego, CA) using unpaired two-tailed Student’s *t* test. Significance was established at P-values < 0.05.

## Acknowledgements

We thank J. Casadesús (University of Seville, Spain) and M. Hensel (University of Osnabrück, Germany) for the gift of *S.* Typhimurium strains and, L.A. Fernández (CNB-CSIC, Madrid, Spain) for the expression vector derivate of vector pGEN222 used for the construction of the reporter plasmids. We also thank Gadea Rico-Peréz for the construction of some tagged S. Typhimurium strains and Henar González for the technical support. D.L.-E. is supported by a PhD fellowship from the “Programa de Formación de Personal Investigador (FPI)” of the Spanish Ministry of Science and Innovation (ref. BES-2017-080709). This work was supported by grant PID2020-112971GB-I00 from the Spanish Ministry of Science and Innovation to F.G-dP.

---

**Table S1.** Bacterial strains and plasmids used in the study

| Bacterial strain/<br>plasmid | Relevant genotype | Source/<br>reference |
| --- | --- | --- |
| S. Typhimurium |  |  |
| SV5015 | SL1344, *hisG*<sup>+</sup> | (Vivero et al., 2008) |
| MD5064 | SV5015 PBP2<sub>SAL</sub>-3xFLAG PBP3<sub>SAL</sub>-3xFLAG | (Castanheira et al., 2020) |
| MD5080 | SV5015 *ompR1009::Tn10* PBP2<sub>SAL</sub>-3xFLAG PBP3<sub>SAL</sub>-3xFLAG | This study |
| MD5540 | SV5015 *phoP7953::Tn10* PBP2<sub>SAL</sub>-3xFLAG PBP3<sub>SAL</sub>-3xFLAG | This study |
| MD2540 | SV5015 PBP2<sub>SAL</sub>-3xFLAG::Km<sup>R</sup> | This study |
| MD3841 | SV5015 *ompR1009::Tn10* PBP2<sub>SAL</sub>-3xFLAG | This study |
| MD3896 | SV5015 *phoP7953::Tn10* PBP2<sub>SAL</sub>-3xFLAG | This study |
| MD3894 | SV5015 *slyA::pRR10ΔtrfAPen<sup>R</sup>* PBP2<sub>SAL</sub>-3xFLAG::Km<sup>R</sup> | This study |
| MD5140 | SV5015 *ssrB::Km<sup>R</sup>* PBP2<sub>SAL</sub>-3xFLAG | M. Hensel |
| MD2559 | SV5015 PBP3<sub>SAL</sub>-3xFLAG | (Castanheira et al., 2017) |
| MD3842 | SV5015 *ompR1009::Tn10* PBP3<sub>SAL</sub>-3xFLAG | (Castanheira et al., 2017) |
| MD3897 | SV5015 *phoP7953::Tn10* PBP3<sub>SAL</sub>-3xFLAG | (Castanheira et al., 2017) |
| MD3895 | SV5015 *slyA::pRR10ΔtrfAPen<sup>R</sup>* PBP3<sub>SAL</sub>-3xFLAG::Km<sup>R</sup> | (Castanheira et al., 2017) |
| MD5139 | SV5015 *ssrB::Km<sup>R</sup>* PBP3<sub>SAL</sub>-3xFLAG | M. Hensel |
| MD5416 | SV5015 *Δprc::Km<sup>R</sup>* PBP2<sub>SAL</sub>-3xFLAG PBP3<sub>SAL</sub>-3xFLAG | This study |
| MD5420 | SV5015 *ompR1009::Tn10 Δprc::Km<sup>R</sup>* PBP2<sub>SAL</sub>-3xFLAG PBP3<sub>SAL</sub>-3xFLAG | This study |
| MD5421 | SV5015 *mrcA-3xFLAG::Km<sup>R</sup>* PBP2<sub>SAL</sub>-3xFLAG PBP3<sub>SAL</sub>-3xFLAG | This study |
| MD5422 | SV5015 *mrcB-3xFLAG::Km<sup>R</sup>* PBP2<sub>SAL</sub>-3xFLAG PBP3<sub>SAL</sub>-3xFLAG | This study |
| MD5425 | SV5015 pAC-6xHis-*ftsI* (6xHis-PBP3) | This study |
| MD5426 | SV5015 *Δprc::Km<sup>R</sup>* pAC-6xHis-*ftsI* (6xHis-PBP3) | This study |
| MD5427 | SV5015 pAC-6xHis-PBP3<sub>SAL</sub> | This study |
| MD5428 | SV5015 *Δprc::Km<sup>R</sup>* pAC-6xHis-PBP3<sub>SAL</sub> | This study |
| MD5436 | SV5015 pGEN222(Δ*PompC-ova*)-PPBP2<sub>SAL</sub>-*gfp*<sup>TCD</sup> | This study |
| MD5437 | SV5015 pGEN222(Δ*PompC-ova*)-PPBP3<sub>SAL</sub>-*gfp*<sup>TCD</sup> | This study |
| MD5438 | SV5015 *ompR1009::Tn10* pGEN222(Δ*PompC-ova*)-PPBP2<sub>SAL</sub>-*gfp*<sup>TCD</sup> | This study |
| MD5439 | SV5015 *ompR1009::Tn10* pGEN222(ΔPompC-*ova*)-PPBP3<sub>SAL</sub>-*gfp*<sup>TCD</sup> | This study |
| MD5440 | SV5015 *phoP7953::Tn10* pGEN222(ΔPompC-*ova*)-PPBP2<sub>SAL</sub>-*gfp*<sup>TCD</sup> | This study |
| MD5441 | SV5015 *phoP7953::Tn10* pGEN222(ΔPompC-*ova*)-PPBP3<sub>SAL</sub>-*gfp*<sup>TCD</sup> | This study |
| MD5446 | SV5015 pGEN222(ΔPompC-*ova*)-mut-PPBP2<sub>SAL</sub> (TGTAACAA>GCTCGGAC)- *gfp*<sup>TCD</sup> | This study |
| MD5447 | SV5015 pGEN222(ΔPompC-*ova*)-mut-PPBP3<sub>SAL</sub> (TGTTGCAA>GCTCGGAC)- *gfp*<sup>TCD</sup> | This study |
| MD5449 | SV5015 pGEN222(ΔPompC-*ova*) (promoter-less vector) | This study |
| MD5451 | SV5015 *rpoS::Cm<sup>R</sup>* PBP2<sub>SAL</sub>-3xFLAG PBP3<sub>SAL</sub>-3xFLAG | This study |
| MD5077 | SV5015 *rpoE::Km<sup>R</sup>* PBP2<sub>SAL</sub>-3xFLAG PBP3<sub>SAL</sub>-3xFLAG | This study |
| MD3238 | SV5015 *dacC*-3xFLAG::*Km<sup>R</sup>* | This study |
| MD3843 | SV5015 *ompR1009::Tn10* *dacC*-3xFLAG::*Km<sup>R</sup>* | This study |
| *E. coli* |  |  |
| MG1655 | K-12, wild type |  |
| MD5442 | MG1655 pGEN222(ΔPompC-*ova*)::P-PBP2<sub>SAL</sub>-*gfp*<sup>TCD</sup> | This study |
| MD5443 | MG1655 pGEN222(ΔPompC-*ova*)::P-PBP3<sub>SAL</sub>- *gfp*<sup>TCD</sup> | This study |
| MD5450 | MG1655 pGEN222(ΔPompC-*ova*) (promoter-less vector) | This study |
| Plasmids |  |  |
| pAC-6xHIS-Plac | *Cm<sup>R</sup>* | (Castanheira et al., 2017) |
| pCP20 | FLP+, Amp<sup>R</sup>, *Cm<sup>R</sup>* | (Cherepanov and Wackernagel, 1995) |
| pSUB11 | 3xFLAG sequence, *Km<sup>R</sup>* | (Uzzau et al., 2001) |
| pKD46 | $\gamma$, $\beta$, *exo*. Amp<sup>R</sup> | (Datsenko and Wanner, 2000) |
| pGEN222 | Amp<sup>R</sup>; p15A-ori, *hok/sok* stability system, *PompC::gfp*-UV | (Galen et al., 1999) |
| pGEN222-*ova-gfp* | pGEN222-PompC-*ova-gfp*<sup>TCD</sup> | L.A. Fernández |

**Table S2.**
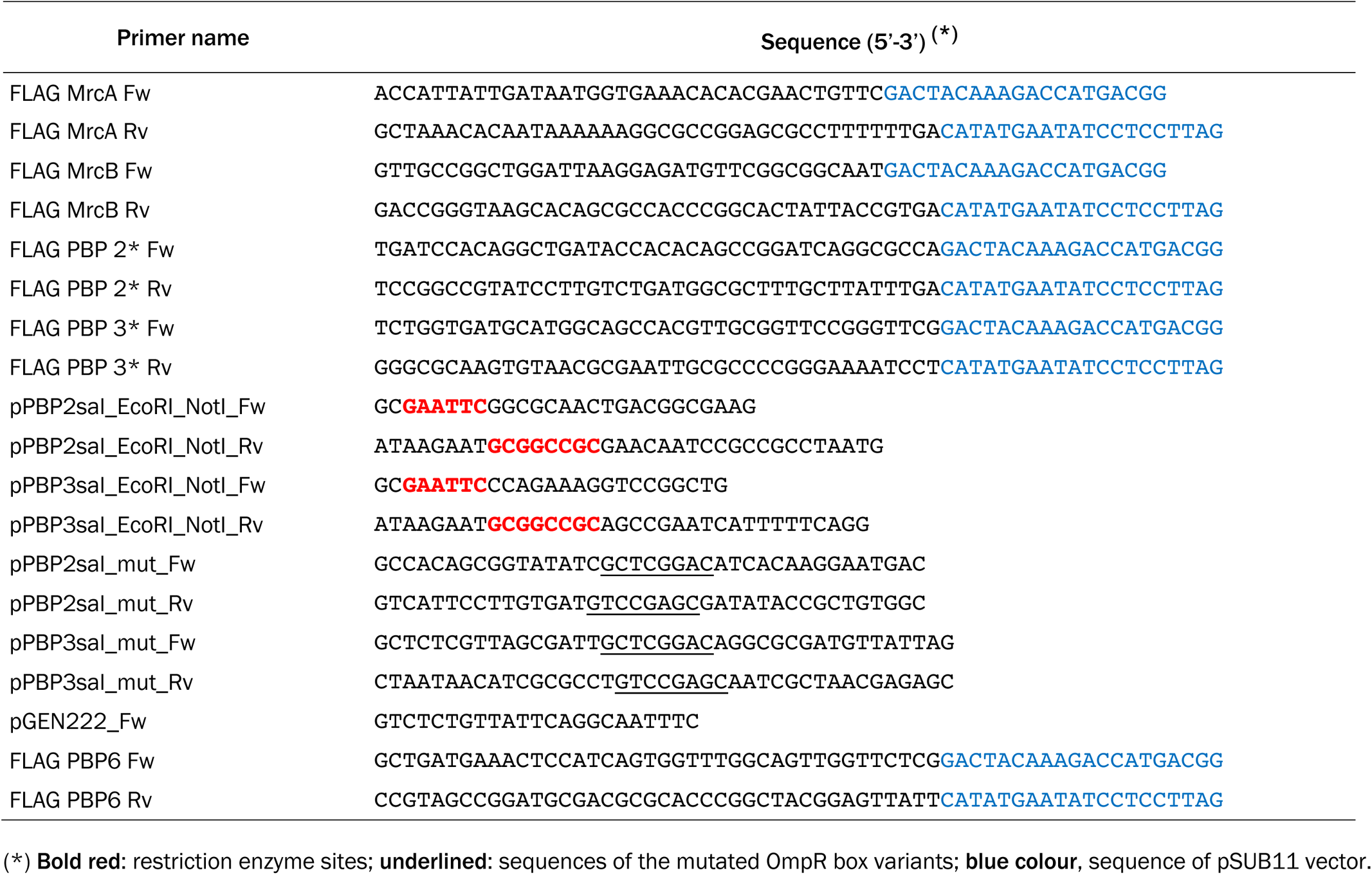
Oligonucleotides used in this study as primers

## Notes

### Competing Interest Statement

The authors have declared no competing interest.

